## Supplementary table S1 and figures S1-3 for "A socio-ecological System Dynamics model of antimicrobial use and resistance"

### Supplementary material

- Model files with documentation
  - AMR_LTG_anecdotal.stm (model from Fig. 3a)
  - AMR_LTG_surveillance.stm (model from Fig. 3b)
  - These Stella Architect files can be opened with isee Player (free software): <https://www.iseesystems.com/store/products/player.aspx>
- Sensitivity run outputs (.xls file)
- Settings used for sensitivity analysis (Table S1)
- Additional sensitivity run figures (Figures S1-S3)

Table S1. Settings used for sensitivity analysis

| **Parameter(s) assessed in sensitivity analysis** | **Parameter range explored** | **Values of other parameters** | **Sensitivity analysis settings** |
| --- | --- | --- | --- |
| a) timescale of microbial evolution and b) time horizon of clinician judgment (latter tested in anecdotal version only) | 0.1-26 weeks for timescale of microbial evolution  1-26 weeks for time horizon of clinician judgment (in anecdotal version only) | Initial susceptible fraction 1, relative fitness in presence of antimicrobial 0.95, relative fitness in absence of antimicrobial 0.05, mean antimicrobial course length 2 weeks | Limited runs (Sobol sequencing); 50 runs; run duration 2080 weeks for anecdotal prescribing version; 520 weeks for surveillance variant. Additional 25 runs with timescale of microbial evolution at 0.1-2 weeks for surveillance. |
| Relative fitness of resistant form in a) presence and b) absence of antimicrobial | 1×10^-12^ to 1 for relative fitness under both conditions | Initial susceptible fraction 1, time horizon of clinician judgment 10 weeks (anecdotal prescribing variant only), timescale of microbial evolution 5 weeks, mean antimicrobial course length 2 weeks | Limited runs (Sobol sequencing); 50 runs; run duration 2080 weeks for anecdotal prescribing version; 520 weeks for surveillance. |

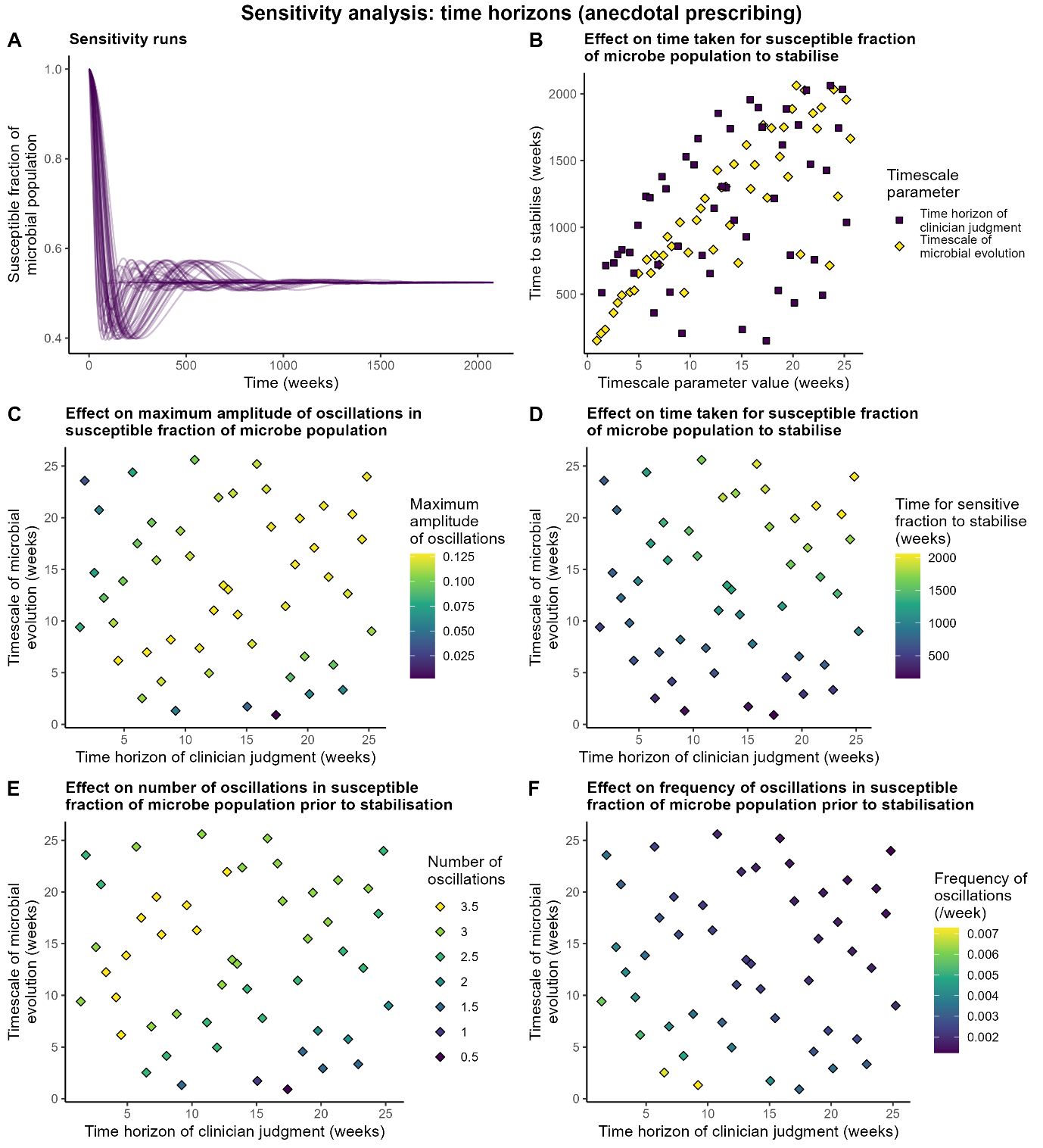

Figure S1. Sensitivity analysis findings: time horizon parameters (anecdotal prescribing)

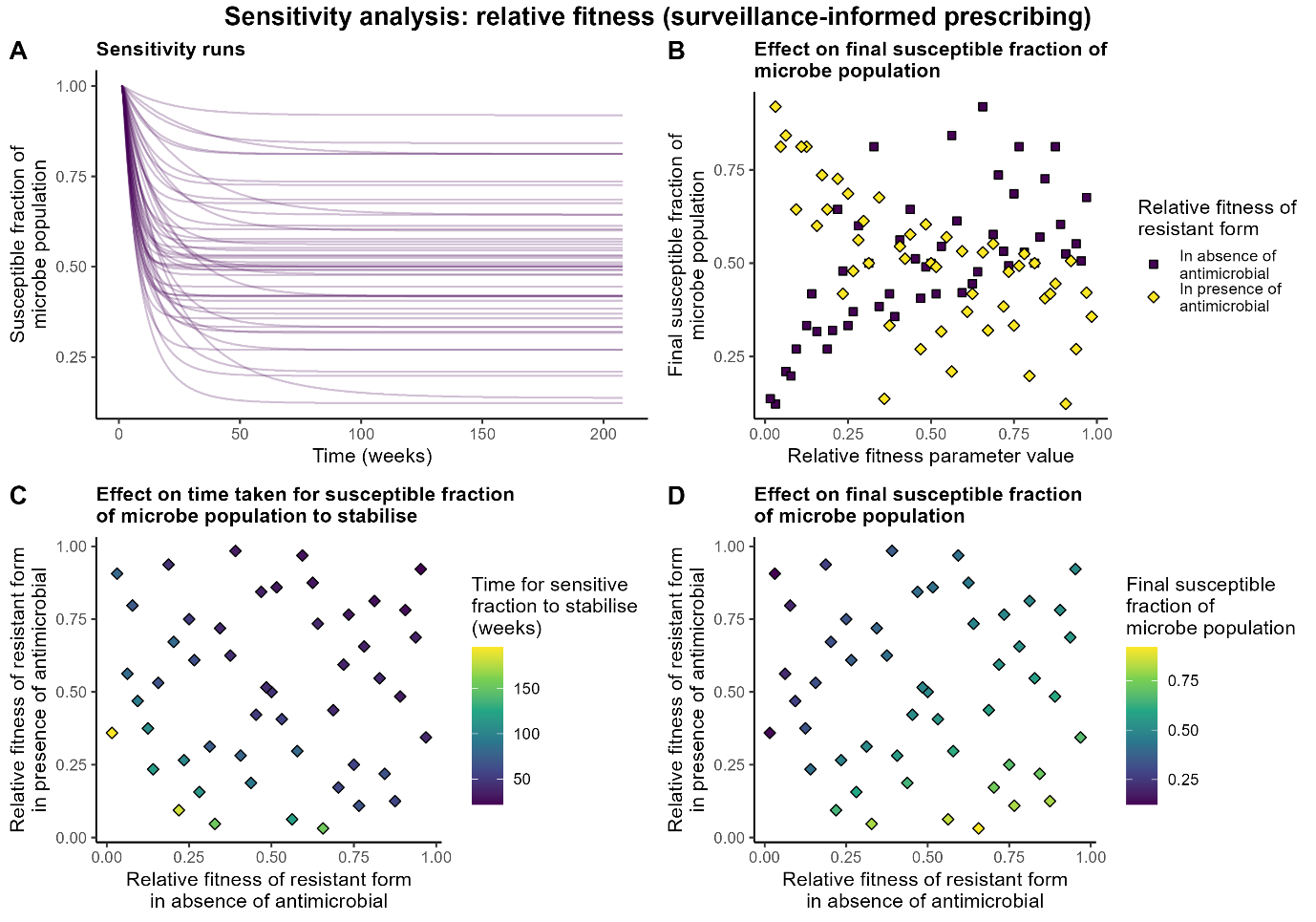

*Figure S2. Sensitivity analysis findings: relative fitness parameters (surveillance-informed prescribing)*

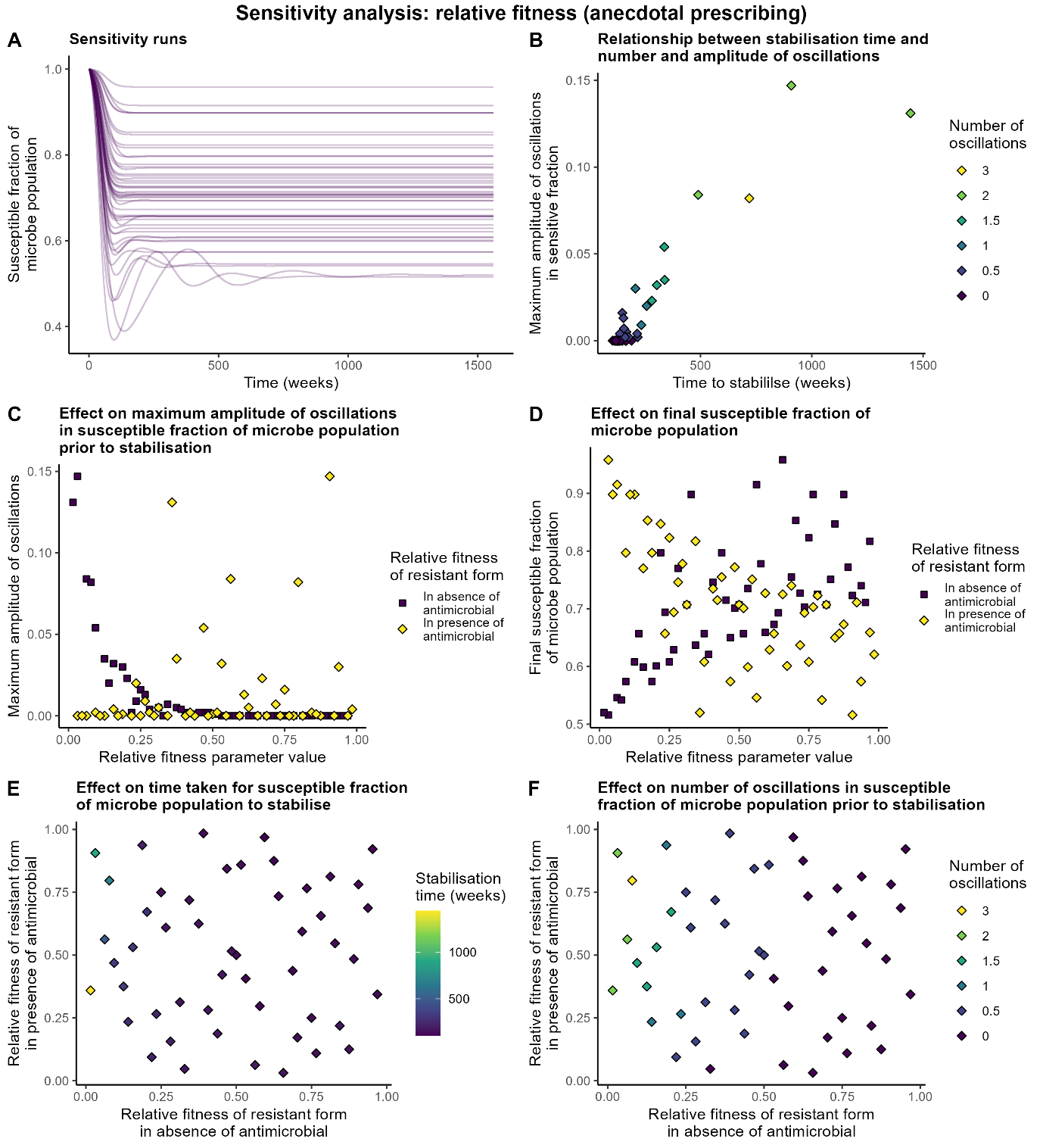

Figure S3. Sensitivity analysis findings: relative fitness parameters (anecdotal prescribing)
